# Explicit Ion Modeling Predicts Physicochemical Interactions for Chromatin Organization

**DOI:** 10.1101/2023.05.16.541030

**Authors:** Xingcheng Lin, Bin Zhang

## Abstract

Molecular mechanisms that dictate chromatin organization *in vivo* are under active investigation, and the extent to which intrinsic interactions contribute to this process remains debatable. A central quantity for evaluating their contribution is the strength of nucleosome-nucleosome binding, which previous experiments have estimated to range from 2 to 14 ***k_B_T***. We introduce an explicit ion model to dramatically enhance the accuracy of residue-level coarse-grained modeling approaches across a wide range of ionic concentrations. This model allows for *de novo* predictions of chromatin organization and remains computationally efficient, enabling large-scale conformational sampling for free energy calculations. It reproduces the energetics of protein-DNA binding and unwinding of single nucleosomal DNA, and resolves the differential impact of mono and divalent ions on chromatin conformations. Moreover, we showed that the model can reconcile various experiments on quantifying nucleosomal interactions, providing an explanation for the large discrepancy between existing estimations. We predict the interaction strength at physiological conditions to be 9 ***k_B_T***, a value that is nonetheless sensitive to DNA linker length and the presence of linker histones. Our study strongly supports the contribution of physicochemical interactions to the phase behavior of chromatin aggregates and chromatin organization inside the nucleus.

## Introduction

Three-dimensional genome organization plays essential roles in numerous DNA-templated processes.^1–5^ Understanding the molecular mechanisms for its establishment could improve our understanding of these processes and facilitate genome engineering. Advancements in high-throughput sequencing and microscopic imaging have enabled genome-wide structural characterization, revealing a striking compartmentalization of chromatin at large scales.^6–9^ For example, A compartments are enriched with euchromatin and activating post-translational modifications to histone proteins. They are often spatially segregated from B compartments that enclose heterochromatin with silencing histone marks.^3,4,10–12^

Compartmentalization has been proposed to arise from the microphase separation of different chromatin types as in block copolymer systems.^13–25^ However, the molecular mechanisms that drive the microphase separation are not yet fully understood. Protein molecules that recognize specific histone modifications have frequently been found to undergo liquid-liquid phase separation,^23,26–31^ potentially contributing to chromatin demixing. Demixing can also arise from interactions between chromatin and various nuclear landmarks such as nuclear lamina and speckles,^11,15,25,32^ as well as active transcriptional processes.^33–36^ Furthermore, recent studies have revealed that nucleosome arrays alone can undergo spontaneous phase separation,^37–39^ indicating that compartmentalization may be an intrinsic property of chromatin driven by nucleosome-nucleosome interactions.

The relevance of physicochemical interactions between nucleosomes to chromatin organization *in vivo* has been constantly debated, partly due to the uncertainty in their strength.^40–43^ Examining the interactions between native nucleosomes poses challenges due to the intricate chemical modifications that histone proteins undergo within the nucleus and the variations in their underlying DNA sequences.^44,45^ Many *in vitro* experiments have opted for reconstituted nucleosomes that lack histone modifications and feature well-positioned 601-sequence DNA to simplify the chemical complexity. These experiments aim to establish a fundamental reference point, a baseline for understanding the strength of interactions within native nucleosomes. Nevertheless, even with reconstituted nucleosomes, a consensus regarding the significance of their interactions remains elusive. For example, using force-measuring magnetic tweezers, Kruithof et al. estimated the inter-nucleosome binding energy to be *∼* 14 k_B_T.^40^ On the other hand, Funke et al. introduced a DNA origami-based force spectrometer to directly probe the interaction between a pair of nucleosomes,^43^ circumventing any potential complications from interpretations of single molecule traces of nucleosome arrays. Their measurement reported a much weaker binding free energy of approximately 2 k_B_T. This large discrepancy in the reported reference values complicates a further assessment of the interactions between native nucleosomes and their contribution to chromatin organization *in vivo*.

Computational modeling is well suited for reconciling the discrepancy across experiments and determining the strength of internucleosome interactions. The high computational cost of atomistic simulations^46–48^ have inspired several groups to calculate the nucleosome binding free energy with coarse-grained models.^49,50^ However, the complex distribution of charged amino acids and nucleotides at nucleosome interfaces places a high demand on force field accuracy. In particular, most existing models adopt a mean-field approximation with the Debye-Hückel theory^51^ to describe electrostatic interactions in an implicit-solvent environment,^49,50,52,53^ preventing an accurate treatment of the complex salt conditions explored in experiments. Further force field development is needed to improve the accuracy of coarsegrained modeling across different experimental settings.^54–57^

We introduce a residue-level coarse-grained explicit-ion model for simulating chromatin conformations and quantifying inter-nucleosome interactions. We validate our model’s accuracy through extensive simulations, demonstrating that it reproduces the binding affinities of protein-DNA complexes ^58^ and energetic cost of nucleosomal DNA unwinding.^59^ Further simulations of chromatin at various salt concentrations reproduce experimentally measured sedimentation coefficients.^60^ We also reveal extensive close contacts between histone proteins and DNA across nucleosomes, the perturbation of which explains the discrepancy among various experimental studies. Finally, we determined the binding free energy between a pair of nucleosomes under physiological salt concentrations as *∼* 9 k*_B_*T. While longer linker DNA would reduce this binding energy, linker histones can more than compensate this reduction to mediate inter-nucleosome interactions with disordered, charged terminal tails. Our study supports the importance of intrinsic physicochemical interactions in chromatin organization *in vivo*.

## Results

### Counterion condensation accommodates nucleosomal DNA unwrapping

Various single-molecule studies have been carried out to probe the stability of nucleosomes and the interactions between histone proteins and DNA. ^41,59,61–63^ The DNA-unzipping experiment performed by Hall et al.^59^ is particularly relevant since the measured forces can be converted into a free energy profile of DNA unwinding at a base-pair resolution, as shown by Forties et al. with a continuous-time Markov model.^64^ The high-resolution quantification of nucleosome energetics is valuable for benchmarking the accuracy of computational models. We introduce a coarse-grained explicit-ion model for chromatin simulations (Fig. 1).

**Figure 1:**
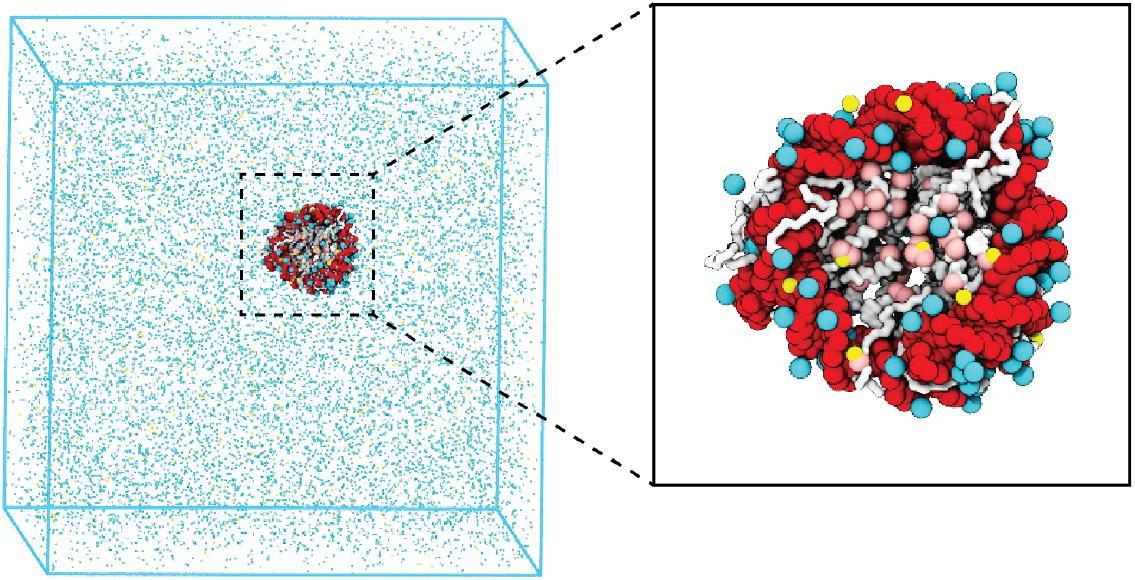
Illustration of the residue-level coarse-grained explicit-ion model for chromatin simulations. The left panel presents a snapshot for the simulation box of a 147-bp nucleosome in a solution of 100 mM NaCl and 0.5 mM MgCl_2_. The nucleosomal DNA and histone proteins are colored in red and white, respectively. The Zoom-in on the right highlights the condensation of ions around the nucleosome, with Na^+^ in cyan and Mg^2+^ in yellow. Negative residues of the histone proteins are colored in pink.

The model represents each amino acid with one coarse-grained bead and three beads per nucleotide. It resolves the differences among various chemical groups to accurately describe biomolecular interactions with physical chemistry potentials. Our explicit representation of monovalent and divalent ions enables a faithful description of counter ion condensation and its impact on electrostatic interactions between protein and DNA molecules. Additional model details are provided in the *Materials and Methods* and the Supporting Information.

We performed umbrella simulations^65^ to determine the free energy profile of nucleosomal DNA unwinding. The experimental buffer condition of 0.10M NaCl and 0.5mM MgCl_2_ ^59^ was adopted in simulations for direct comparison. As shown in Fig. 2B, the simulated values match well with experimental results over a wide range. Furthermore, we computed the binding free energy for a diverse set of protein-DNA complexes and the simulated values again match well with experimental data (Fig. S1), supporting the model’s accuracy.

**Figure 2:**
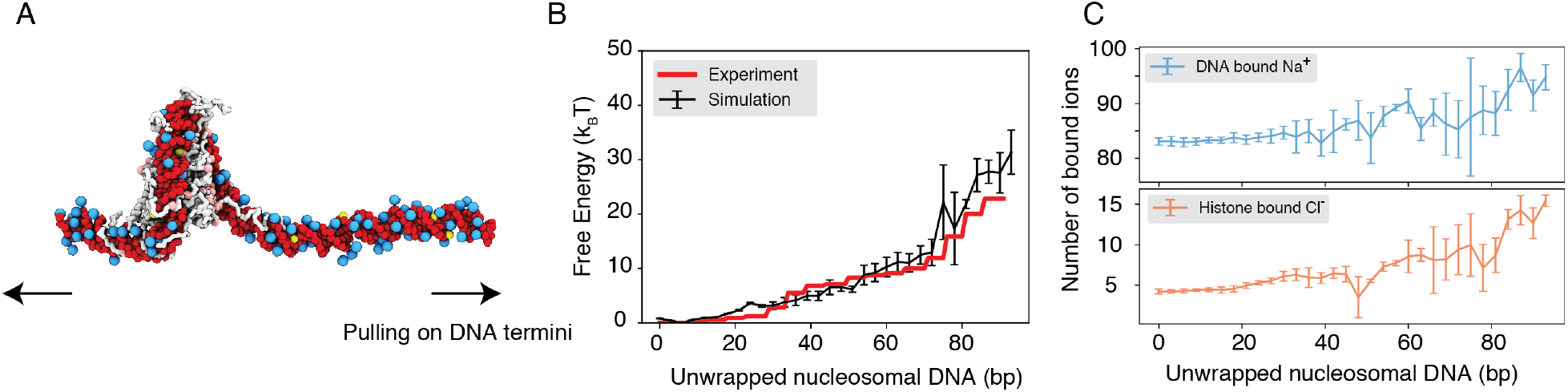
Explicit ion modeling reproduces the energetics of nucleosomal DNA unwrapping. (A) Illustration of the umbrella simulation setup using the end-to-end distance between two DNA termini as the collective variable. The same color scheme as in Fig. 1 is adopted. Only ions close to the nucleosomes are shown for clarity. (B) Comparison between simulated (black) and experimental (red) free energy profile as a function of the unwrapped DNA base pairs. Error bars were computed as the standard deviation of three independent estimates. (C) The average number of Na^+^ ions within 10 ^°^A of the nucleosomal DNA (top) and Cl*^−^* ions within 10 ^°^A of histone proteins (bottom) are shown as a function of the unwrapped DNA base pairs.

Counterions are often released upon protein-DNA binding to make room for close contacts at the interface, contributing favorably to the binding free energy in the form of entropic gains.^66^ However, previous studies have shown that the histone-DNA interface in a fully wrapped nucleosome configuration is not tightly sealed but instead permeated with water molecules and mobile ions. ^67,68^ Given their presence in the bound form, how these counterions contribute to nucleosomal DNA unwrapping remains to be shown. We calculated the number of DNA-bound cations and protein-bound anions as DNA unwraps. Our results, shown in Fig. 2C, indicate that only a modest amount of extra Na^+^ and Cl*^−^* ions becomes associated with the nucleosome as the outer DNA layer unwraps. However, significantly more ions become bound when the inner layer starts to unwrap (after 73 bp). These findings suggest that counterion release may contribute more significantly to the inner layer wrapping, potentially caused by a tighter protein-DNA interface.

### Charge neutralization with *Mg*^2+^ compacts chromatin

In addition to contributing to the stability of individual nucleosomes, counterions can also impact higher-order chromatin organization. Numerous groups have characterized the structures of nucleosome arrays, ^60,69–73^ revealing a strong dependence of chromatin folding on the concentration and valence of cations.

To further understand the role of counterions in chromatin organization, we studied a 12-mer with 20-bp-long linker DNA under different salt conditions. We followed the experiment setup by Correll et al.^60^ that immerses chromatin in solutions with 5mM NaCl, 150mM NaCl, 0.6mM MgCl_2_, or 1mM MgCl_2_. To facilitate conformational sampling, we carried out umbrella simulations with a collective variable that quantifies the similarity between a given configuration and a reference two-start helical structure. Simulation details and the precise definition of the collective variable are provided in Supporting Information. Data from different umbrella windows were combined together with proper reweighting^74^ for analysis. As shown in Fig. 3A, the average sedimentation coefficients determined from our simulations match well with experimental values. Specifically, the simulations reproduce the strong contrast in chromatin size between the two systems with different NaCl concentrations. Chromatin under 5 mM NaCl features an extended configuration with minimal stacking between 1-3 nucleosomes (Fig. 3B). On the other hand, the compaction is evident at 150 mM NaCl. Notably, in agreement with previous studies,^75–78^ we observe tri-nucleosome configurations as chromatin extends. Finally, the simulations also support that divalent ions are more effective in packaging chromatin than NaCl. Even in the presence of 0.6 mM MgCl_2_, the chromatin sedimentation coefficient is comparable to that obtained at 150 mM of NaCl.

**Figure 3:**
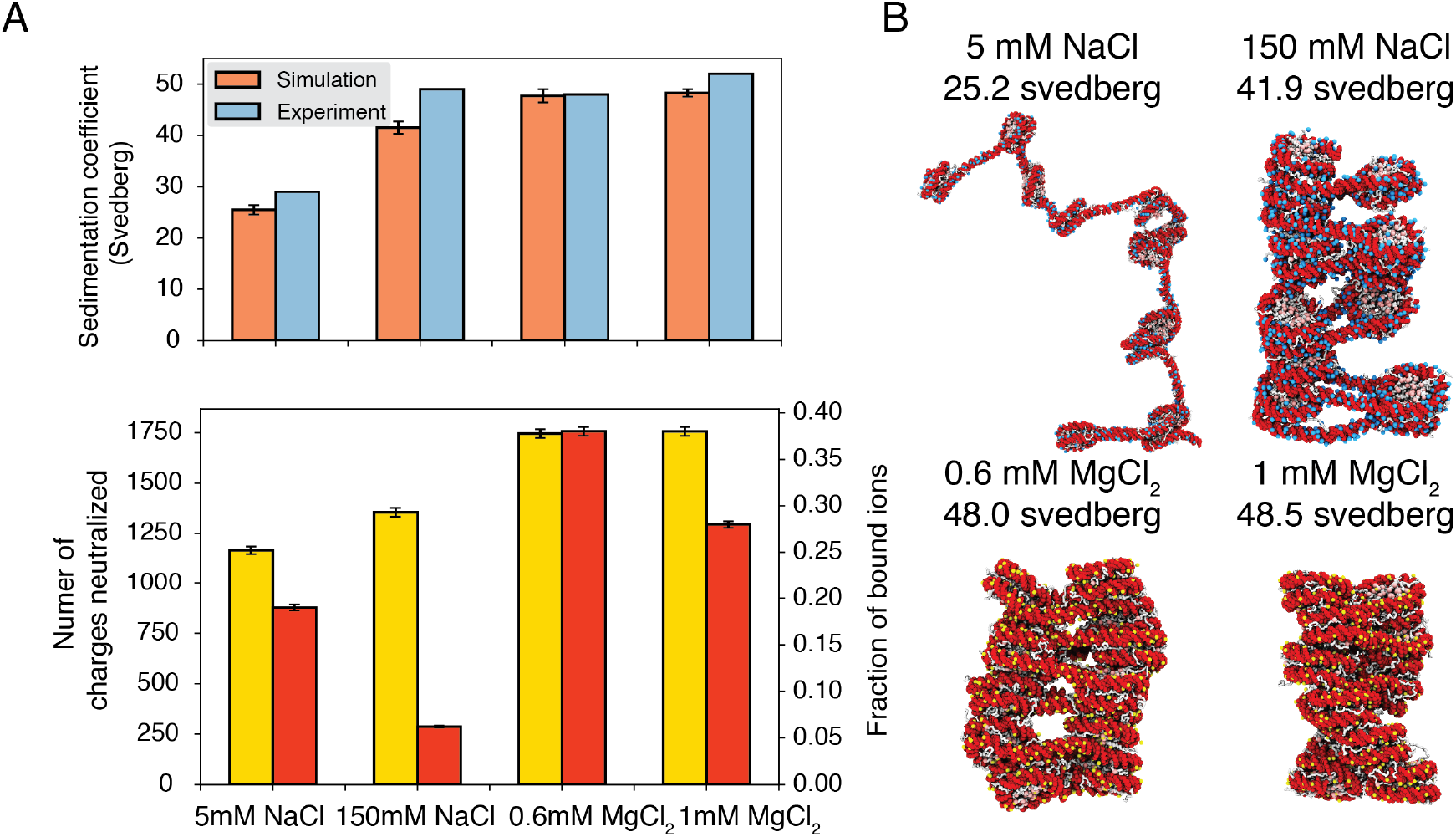
Explicit ion modeling predicts salt-dependent conformations of a 12-mer nucleosome array. (A) Top: Comparison of simulated and experimental^60^ sedimentation coefficients of chromatin at different salt concentrations. Bottom: Number of DNA charges neutralized by bound cations (yellow, left y-axis label) and the fraction of ions bound to DNA (red, right y-axis label) at different salt concentrations. The error bars were estimated from the standard deviation of simulated probability distributions (Fig. S2) (B) Representative chromatin structures with sedimentation coefficients around the mean values at different salt concentrations.

We further characterized ions that are in close contact with DNA to understand their impact on chromatin organization. Our simulations support the condensation of cations, especially for divalent ions (Fig. 3A bottom) as predicted by the Manning theory.^79,80^ Ion condensation weakens the repulsion among DNA segments that prevents chromatin from collapsing. Notably, the fraction of bound Mg^2+^ is much higher than Na^+^. Correspondingly, the amount of neutralized negative charges is always greater in systems with divalent ions, despite the significantly lower salt concentrations. The difference between the two types of ions arises from the more favorable interactions between Mg^2+^ and phosphate groups that more effectively offset the entropy loss due to ion condensation.^80^ While higher concentrations of NaCl do not dramatically neutralize more charges, the excess ions provide additional screening to weaken the repulsion among DNA segments, stabilizing chromatin compaction.

### Close contacts drive nucleosome binding free energy

Encouraged by the explicit ion model’s accuracy in reproducing experimental measurements of single nucleosomes and nucleosome arrays, we moved to directly quantify the strength of inter-nucleosomes interactions. We once again focus on reconstituted nucleosomes for a direct comparison with *in vitro experiments*. These experiments have yielded a wide range of values, ranging from 2 to 14 k*_B_*T.^40,41,43^ Accurate quantification will offer a reference value for conceptualizing the significance of physicochemical interactions among native nucleosomes in chromatin organization *in vivo*.

To reconcile the discrepancy among various experimental estimations, we directly calculated the binding free energy between a pair of nucleosomes with umbrella simulations. We adopted the same ionic concentrations as in the experiment performed by Funke et al.^43^ with 35 mM NaCl and 11 mM MgCl_2_. We focus on this study since the experiment directly measured the inter-nucleosomal interactions, allowing straightforward comparison with simulation results. Furthermore, the reported value for nucleosome binding free energy deviates the most from other studies. In one set of umbrella simulations, we closely mimicked the DNA origami device employed by Funke et al. to move nucleosomes along a predefined path for disassociation (Fig. 4A, A1 to A3). For example, neither nucleosome can freely rotate (Fig. S3); the first nucleosome is restricted to the initial position, and the second nucleo-some can only move within the Y-Z plane along the arc 15 nm away from the origin. For comparison, we performed a second set of independent simulations without imposing any restrictions on nucleosome orientations. Additional simulation details can be found in the Supporting Information.

**Figure 4:**
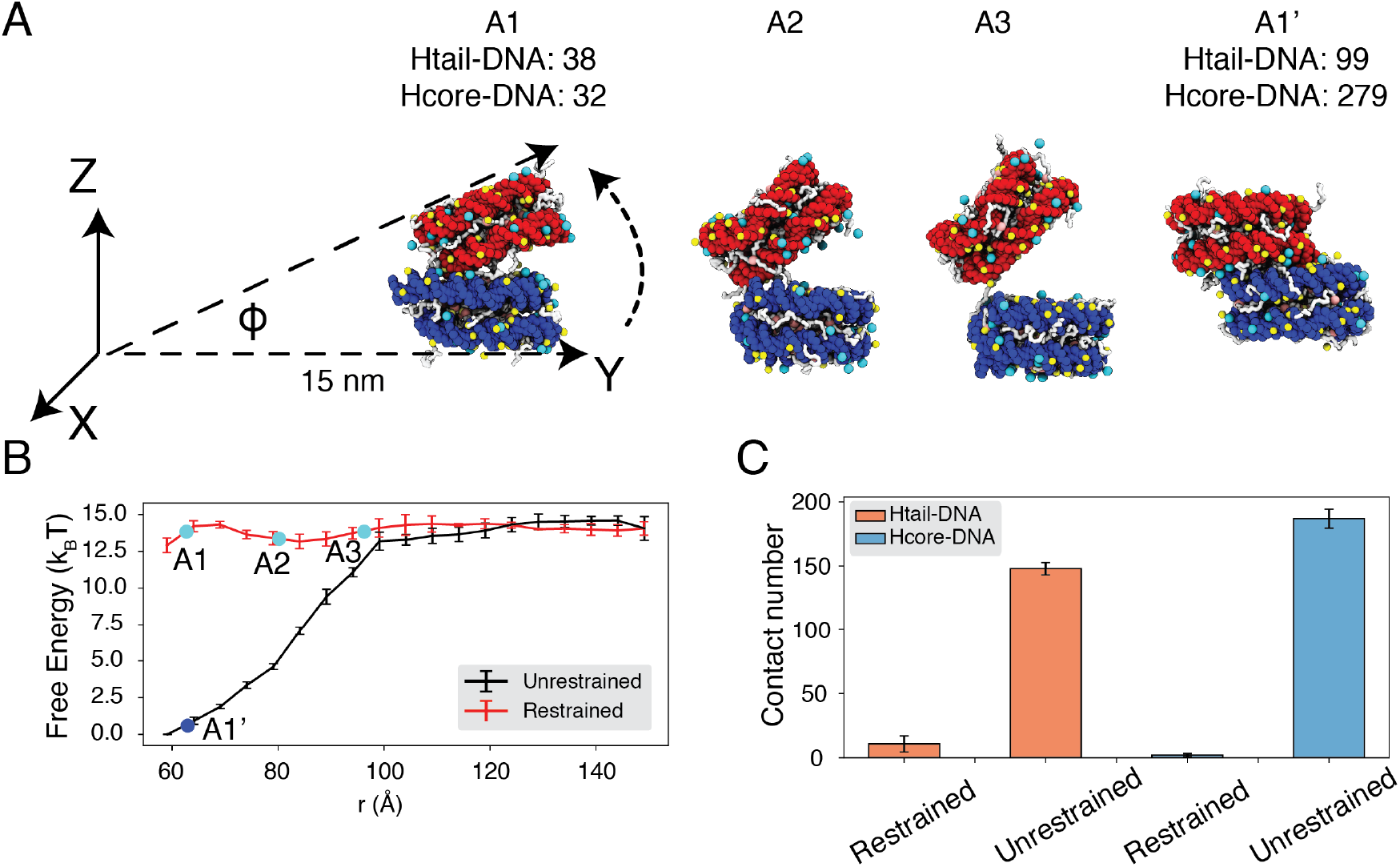
Close contacts give rise to strong internucleosomal interactions. (A) Illustration of the simulation protocol employed to mimic the nucleosome unbinding pathway dictated by the DNA origami device.^43^ The three configurations, A1, A2, and A3, corresponding to the three cyan dots in part B at distances 62.7, 80.2, and 96.3^°^A. For comparison, a tightly bound configuration uncovered in simulations without any restraints of nucleosome movement is shown as A1’. The number of contacts formed by histone tails and DNA (Htail-DNA) and by histone core and DNA (Hcore-DNA) from different nucleosomes are shown for A1 and A1’. (B) Free energy profile as a function of the distance between the geometric centers of the two nucleosomes, computed from unrestrained (black) and DNA origami-restrained simulations (red). Error bars were computed as the standard deviation of three independent estimates. (C) Average inter-nucleosomal contacts between DNA and histone tail (orange) and core (blue) residues, computed from unrestrained and DNA origami-restrained simulations. Error bars were computed as the standard deviation of three independent estimates.

Strikingly, the two sets of simulations produced dramatically different binding free energies. Restricting nucleosome orientations produced a binding free energy of *∼* 2k*_B_*T, reproducing the experimental value (Fig. 4B, S4). On the other hand, the binding free energy increased to 15 k*_B_*T upon removing the constraints. Further examination of inter-nucleosomal contacts revealed the origin of the dramatic difference in nucleosome binding free energies. As shown in Fig. 4C, the average number of contacts formed between histone tails and DNA from different nucleosomes is around 150 and 10 in the two sets of simulations. A similar trend is observed for histone core-DNA contacts across nucleosomes. The differences are most dramatic at small distances (Fig. S5) and are clearly visible in the most stable configurations. For example, from the unrestricted simulations, the most stable binding mode corresponds to a configuration in which the two nucleosomes are almost parallel to each other (see configuration A1’ in Fig. 4A), with the angle between the two nucleosome planes close to zero (Fig. S6C). However, the inherent design of the DNA-origami device renders this binding mode inaccessible, and the smallest angle between the two nucleosome planes is around 23*^◦^* (see configuration A1 in Fig. 4A). Therefore, a significant loss of inter-nucleosomal contacts caused the small binding free energy seen experimentally.

### Modulation of nucleosome binding free energy by *in vivo* factors

The predicted strength for unrestricted inter-nucleosome interactions supports their significant contribution to chromatin organization *in vivo*. However, the salt concentration studied above and in the DNA origami experiment is much higher than the physiological value.^42^ To further evaluate the *in vivo* significance of inter-nucleosome interactions, we computed the binding free energy at the physiological salt concentration of 150 mM NaCl and 2 mM of MgCl_2_.

We observe a strong dependence of nucleosome orientations on the inter-nucleosome distance. A collective variable, θ, was introduced to quantify the angle between the two nucleosomal planes (Fig. 5A). As shown in two-dimensional binding free energy landscape of internucleosome distance, r, and θ (Fig. 5B), at small distances (*∼*60 ^°^A), the two nucleosomes prefer a face-to-face binding mode with small θ values. As the distance increases, the nucleosomes will almost undergo a 90*^◦^* rotation to adopt perpendicular positions. Such orientations allow the nucleosomes to remain in contact and is more energetically favorable. The orientation preference gradually diminishes at large distances once the two nucleosomes are completely detached. Importantly, we observed a strong inter-nucleosomal interaction with two nucleosomes wrapped by 147-bp 601-sequence DNA (*∼* 9 k*_B_*T).

**Figure 5:**
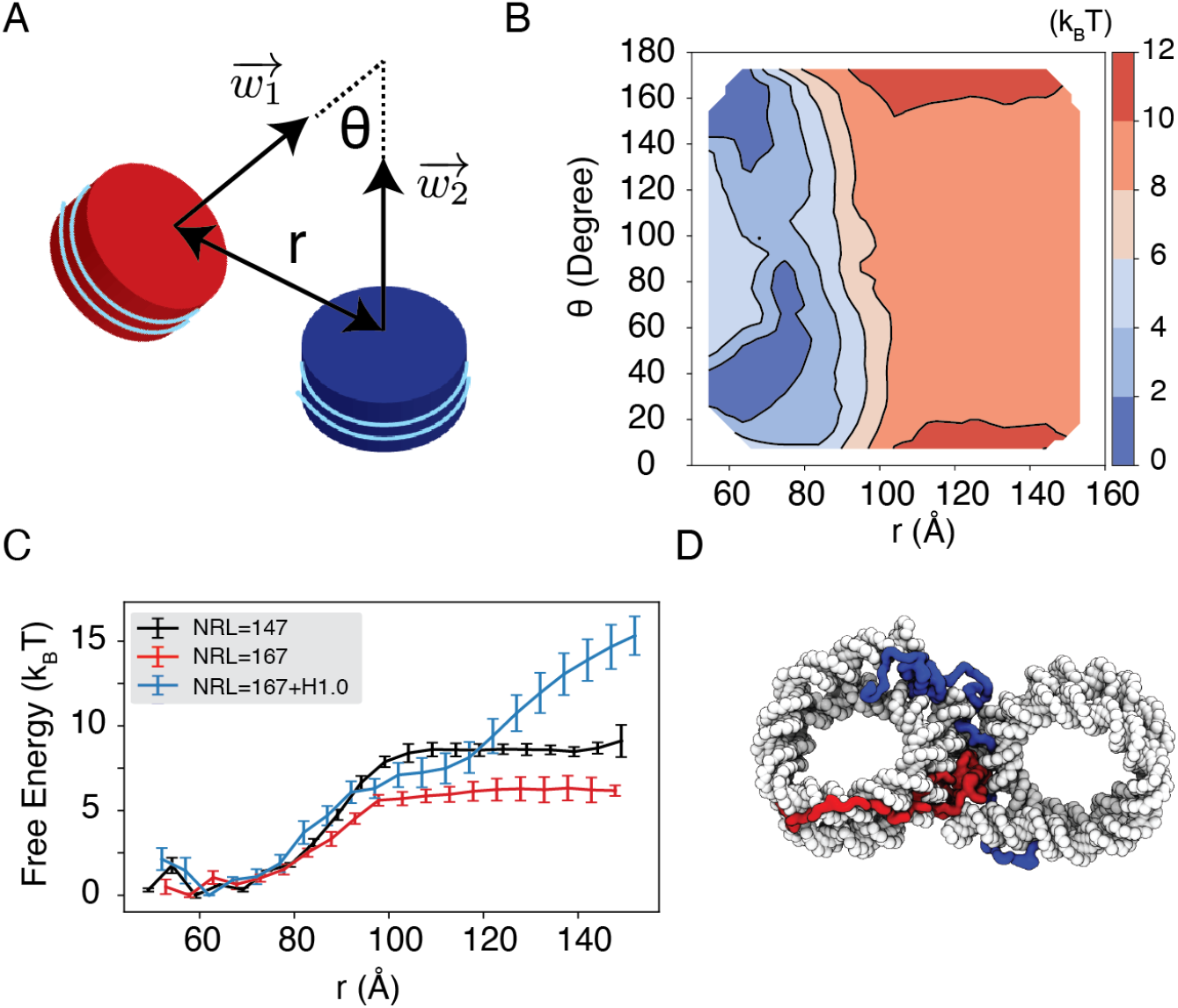
Simulations predict significant internucleosome interactions at physiological conditions. (A) Illustration of the collective variable, θ defined as the angle between two nucleosomal planes, and r defined as the distance between the nucleosome geometric centers. w⃗_1_ and w⃗_2_ represent the axes perpendicular to the nucleosomal planes. (B) The 2D binding free energy profile as a function of θ and r at the physiological salt condition (150mM NaCl and 2mM MgCl_2_) for nucleosomes with the 601 sequence. (C) Dependence of nucleosome binding free energy on nucleosome repeat length (NRL) and linker histone H1.0. (D) Representative structure showing linker histones (orange and green) mediating inter-nucleosomal contacts.

Furthermore, we found that the nucleosome binding free energy is minimally impacted by the precise DNA sequence. For example, when the 601 sequence is replaced with poly-dA:dT or poly-dG:dC, the free energy only varied by *∼*2 k*_B_*T (Fig. S7). However, the poly-dA:dT sequence produced stronger binding while poly-dG:dC weakened the interactions. The sequence specific effects are potentially due to the increased stiffness of poly-dA:dT DNA, ^81^ which causes the DNA to unwrap more frequently, increasing cross nucleosome contacts at larger distances (Fig. S8).

In addition to variations in DNA sequences, *in vivo* nucleosomes also feature different linker lengths. We performed simulations that extend the 601 sequence with 10 extra base pairs of poly-dA:dT sequence at each end, reaching a nucleosome repeat length (NRL) of 167 bp. Consistent with previous studies,^60,82,83^ increasing the NRL weakened inter-nucleosomal interactions (Fig. 5C and Fig. S9), reducing the binding free energy to *∼* 6 k*_B_*T.

Importantly, we found that the weakened interactions upon extending linker DNA can be more than compensated for by the presence of histone H1 proteins. This is demonstrated in Fig. 5C and Fig. S9, where the free energy cost for tearing part two nucleosomes with 167 bp DNA in the presence of linker histones (blue) is significantly higher than the curve for bare nucleosomes (red). Notably, at larger inter-nucleosome distances, the values even exceed those for 147 bp nucleosomes (black). A closer examination of the simulation configurations suggests that the disordered C-terminal tail of linker histones can extend and bind the DNA from the second nucleosome, thereby stabilizing the internucleosomal contacts (as shown in Fig. 5D). Our results are consistent with prior studies that underscore the importance of linker histones in chromatin compaction,^84,85^ particularly in eukaryotic cells with longer linker DNA.^78,86^

## Discussion

We introduced a residue-level coarse-grained model with explicit ions to accurately account for electrostatic contributions to chromatin organization. The model achieves quantitative accuracy in reproducing experimental values for the binding affinity of protein-DNA complexes, the energetics of nucleosomal DNA unwinding, nucleosome binding free energy, and the sedimentation coefficients of nucleosome arrays. It captures the counterion atmosphere around the nucleosome core particle as seen in all-atom simulations^68^ and highlights the contribution of counterions to nucleosome stability. The coarse-grained model also succeeds in resolving the difference between monovalent and divalent ions, supporting the efficacy of divalent ions in neutralizing negative charges and offsetting repulsive interactions among DNA segments.

One significant finding from our study is the predicted strong inter-nucleosome interactions under the physiological salt environment, reaching approximately 9 k*_B_*T. We showed that the much lower value reported in a previous DNA origami experiment is due to the restricted nucleosomal orientation inherent to the device design. Unrestricted nucleosomes allow more close contacts to stabilize binding. A significant nucleosome binding free energy also agrees with the high forces found in single-molecule pulling experiments that are needed for chromatin unfolding.^40,42,87^ We also demonstrate that this strong inter-nucleosomal interaction is largely preserved at longer nucleosome repeat lengths (NRL) in the presence of linker histone proteins. While post-translational modifications of histone proteins may influence inter-nucleosomal interactions, their effects are limited, as indicated by Ding et al. ^75^, and are unlikely to completely abolish the significant interactions reported here. Therefore, we anticipate that, in addition to molecular motors, chromatin regulators, and other molecules inside the nucleus, intrinsic inter-nucleosome interactions are important players in chromatin organization *in vivo*.

We focused our study on single chromatin chains. Strong inter-nucleosome interactions support the compaction and stacking of chromatin, promoting the formation of fibril-like structures. However, as shown in many studies,^39,88–90^ such fibril configurations can hardly be detected *in vivo*. It is worth emphasizing that this lack of fibril configurations does not contradict our conclusion on the significance of inter-nucleosome interactions. In a prior paper, we found that many *in vivo* factors, most notably crowding, could disrupt fibril configurations in favor of inter-chain contacts.^76^ The inter-chain contacts can indeed be driven by favorable inter-nucleosome interactions.

Several aspects of the coarse-grained model presented here can be further improved. For instance, the introduction of specific protein-DNA interactions could help address the differences in non-bonded interactions between amino acids and nucleotides beyond electrostatics.^31^ Such a modification would enhance the model’s accuracy in predicting interactions between chromatin and chromatin-proteins. Additionally, the single-bead-per-amino-acid representation used in this study encounters challenges when attempting to capture the influence of histone modifications, which are known to be prevalent in native nucleosomes. Multiscale simulation approaches may be necessary.^91^ One could first assess the impact of these modifications on the conformation of disordered histone tails using atomistic simulations. By incorporating these conformational changes into the coarse-grained model, systematic investigations of histone modifications on nucleosome interactions and chromatin organization can be conducted. Such a strategy may eventually enable the direct quantification of interactions among native nucleosomes and even the prediction of chromatin organization *in vivo*.

## Methods and Materials

### Coarse-grained modeling of chromatin

The large system size of chromatin and the slow timescale for its conformational relaxation necessitates coarse-grained modeling. Following previous studies,^5,31,75,76,92^ we adopted a residue-level coarse-grained model for efficient simulations of chromatin. The structure-based model^93,94^ was applied to represent the histone proteins with one bead per amino acid and to preserve the tertiary structure of the folded regions. The disordered histone tails were kept flexible without tertiary structure biases. A sequence-specific potential, in the form of the Lennard Jones (LJ) potential and with the strength determined from the Miyazwa-Jernigan (MJ) potential,^95^ was added to describe the interactions between amino acids. The 3SPN.2C model was adopted to represent each nucleotide with three beads and interactions among DNA beads follow the potential outlined in Ref. 96, except that the charge of each phosphate site was switched from −0.6 to −1.0 to account for the presence of explicit ions. The Coulombic potential was applied between charged protein and DNA particles. In addition, a weak, non-specific LJ potential was used to account for the excluded volume effect among all protein-DNA beads. Detail expressions for protein-protein and protein-DNA interaction potentials can be found in Ref. 75 and the Supporting Information.

We observe that residue-level coarse-grained models have been extensively utilized in prior studies to examine the free energy penalty associated with nucleosomal DNA unwinding,^97–99^ sequence-dependent nucleosome sliding,^100,101^ binding free energy between two nucleosomes,^49^ chromatin folding,^75,76^ the impact of histone modifications on tri-nucleosome structures,^102^ and protein-chromatin interactions.^92,103^ The frequent quantitative agreement between simulation and experimental results supports the utility of such models in chromatin studies. Our introduction of explicit ions, as detailed below, further extends the applicability of these models to explore the dependence of chromatin conformations on salt concentrations.

### Coarse-grained modeling of counter ions

Explicit particle-based representations for monovalent and divalent ions are needed to accurately account for electrostatic interactions.^54–57,104–106^ We followed Freeman et al.^54^ to introduce explicit ions (see Fig. 1) and adopted their potentials to describe the interactions between ions and nucleotide particles, with detailed expressions provided in the Supporting Information. Parameters in these potentials were tuned by Freeman et al. ^54^ to reproduce the radial distribution functions and the potential of mean force between ion pairs determined from all-atom simulations.

This explicit ion model was originally introduced for nucleic acid simulations. We generalized the model for protein simulations by approximating the interactions between charged amino acids and ions with parameters tuned for phosphate sites. Parameter values for ion-amino acid interactions are provided in Table S1 and S2.

### Details of molecular dynamics simulations

We simulated various chromatin systems, including a single nucleosome, two nucleosomes, and a 12-mer nucleosome array. The initial configurations for the molecular dynamics simulations were constructed based on the crystal structure of a single nucleosome with PDB ID: 1KX5^67^ and 3LZ1,^107^ or a tetranucleosome with PDB ID: 1ZBB.^108^ We used the 3DNA software^109^ to add additional DNA, connect and align nucleosomes, and extend the chain length as necessary. Further details on constructing the initial configurations are provided in the Supporting Information. Chromatin was positioned at the center of a cubic box with a length selected to avoid interactions between nucleosomes and their periodic images. Counterions were added on a uniformly spaced grid to achieve the desired salt concentrations and to neutralize the system. The number of ions and the size of simulation boxes are provided in Table S3.

All simulations were performed at constant temperature and constant volume (NVT) using the software package LAMMPS.^110^ The electrostatic interactions were implemented with the particle-particle particle-mesh solver, with the relative root-mean-square error in per-atom force set to 0.0001.^111^ A Nośe-Hoover style algorithm^112^ was used to maintain the system temperature at 300K with a damping parameter of 1 ps. We further modeled the histone core and the inner layer of the nucleosomal DNA together as a rigid body to improve computational efficiency. This approximation does not affect the thermodynamic properties of chromatin.^75,76^ Umbrella simulations were used to enhance the sampling of the conformational space,^65^ and details of the collective variables employed in these simulations are provided in the Supporting Information. All the results presented in the main text are reweighted from the biased simulations by the weighted histogram algorithm.^74^

## Acknowledgement

This work was supported by the National Institutes of Health (Grant R35GM133580) and the National Science Foundation (Grant MCB-2042362).

## Data availability

Implementation of the residue-level coarse-grained model and examples for setting up and performing simulations can be found on our GitHub repository.

