## Supporting Information for "Explicit Ion Modeling Predicts Physicochemical Interactions for Chromatin Organization"

#### Summary

**Coarse-grained protein-DNA model** **S-3**

**Coarse-grained explicit ion model** **S-4**

**Molecular dynamics simulation details** **S-5**

Binding free energy of protein-DNA complexes . . . . . . . . . . . . . . . . . . . . . S-5

Single nucleosome simulations for DNA unwinding energetics . . . . . . . . . . . S-6

Ionic dependence of the conformation for a 12-mer nucleosomal array . . . . . . . S-7

Binding free energy between nucleosomes . . . . . . . . . . . . . . . . . . . . . . . S-8

Simulations at high salt concentrations . . . . . . . . . . . . . . . . . . . . . S-8

Simulations at the physiological salt concentration . . . . . . . . . . . . . . . S-10

**Details of simulation analysis** **S-11**

Number of ions bound to DNA and histone proteins . . . . . . . . . . . . . . . . . S-11

Number of unwrapped DNA base pairs . . . . . . . . . . . . . . . . . . . . . . . . S-12

Sedimentation coefficients of nucleosome arrays . . . . . . . . . . . . . . . . . . . . S-12

Estimation of error bars . . . . . . . . . . . . . . . . . . . . . . . . . . . . . . . S-13

### Coarse-grained protein-DNA model

The force fields describing protein-protein, protein-DNA, and DNA-DNA interactions followed previous studies.[S1–S4] Detailed expressions for the potential energies can be found in these references. Specifically, for DNA-DNA interactions, we followed the approach described in Ref. S5, and with the parameters updated to the latest version of the DNA model 3SPN.2C.[S1]

Protein-protein interactions include structure-based terms extracted from the initial configuration and generic terms for specific amino-acid interactions. We first generated the bonded and nonbonded structure-based interactions within histone proteins using the Shadow algorithm[S6] implemented by the SMOG software package.[S7] We further scaled the nonbonded interaction strength[S7] by 2.5 to prevent proteins from unfolding at 300 K. Interactions between histone proteins from different nucleosomes were described using the Miyazwa-Jernigan (MJ) potential[S8] scaled by a factor of 0.4. We have shown in our previous studies that the scaled MJ potential gives a balanced modeling of the radius of gyration for both ordered and disordered proteins.[S3]

Protein-DNA interactions include Colombic interactions and the excluded volume effect. Unlike the previous Debye-Hückel treatment of electrostatic interactions in an implicit-solvent environment, we modeled the electrostatics between proteins and DNA using the Coulombic potential

$$U_{\text{elec}} = \frac{1}{4\pi\epsilon_0}\frac{q_i q_j}{\epsilon_0 r}, \tag{S1}$$

where $\epsilon_0 = 78.0$ is the dielectric constant of the bulk solvent. $q_{\text{i}}$ and $q_{\text{j}}$ correspond to the charges of the two particles. The excluded volume effect was modeled using the WCA potential with the following form

$$U_{\text{WCA}} = \begin{cases} 4\epsilon^{\text{PD}}[(\frac{\sigma}{r})^{12} - (\frac{\sigma}{r})^{6}] + \epsilon^{\text{PD}} & r < r_{\text{cut}} \\ 0 & r > r_{\text{cut}}. \end{cases} \tag{S2}$$

The cutoff distance $r_{\mathrm{cut}}$ was set to $2^{\frac{1}{6}}\sigma$, with $\sigma = 5.7$ Å. The interactive strength $\epsilon^{\mathrm{PD}}$ was set as 0.02987572 kcal/mol. More details of this potential can be found in Ref. S2.

### Coarse-grained explicit ion model

Following Freeman et al.,[S5] we adopted three terms to describe interactions between charged particles and ions: the Coulombic potential for electrostatic interactions, the Gaussian potential for the hydration effect, and the Lennard-Jones potential for the excluded volume effect. Thus

$$U = U_{\mathrm{elec}} + U_{\mathrm{hydr}} + U_{\mathrm{LJ}}. \tag{S3}$$

The electrostatic potential

$$U_{\mathrm{elec}} = \frac{1}{4\pi\epsilon_0}\frac{q_i q_j}{\epsilon_{\mathrm{D}}(r)r}, \tag{S4}$$

where $\epsilon_{\mathrm{D}}(r)$ is a distance-dependent dielectric constant given in the form

$$\epsilon_{\mathrm{D}}(r) = (\frac{5.2+\epsilon_s}{2}) + (\frac{5.2+\epsilon_s}{2})\tanh[\frac{\mathrm{r}-\mathrm{r}_{\mathrm{m}\epsilon}}{\sigma_\epsilon}]. \tag{S5}$$

$\epsilon_s = 78.0$ is the dielectric constant of the bulk solvent, and values for $\sigma_\epsilon$ and $r_{m\epsilon}$ are provided in Table S1. The distance cutoff of this potential is set at 20.0 Å. Electrostatic interactions outside this cutoff are computed in reciprocal space.

The hydration potential

$$U_{\mathrm{hydr}} = \frac{H}{\sigma_h\sqrt{2\pi}}\exp[-\frac{(r-r_{mh})^2}{2\sigma_h^2}]. \tag{S6}$$

$r_{mh}$, $\sigma_h$ and $H$ represent the midpoint, the width, and the height of the hydration shell, respectively, and their ion specific values are provided in Table S1. Any pair of ions experience one hydration potential defined above. For pairs formed with ions of distinct types, a second hydration potential with a different set of parameters is applied, with parameters provided in Table S1. The distance cutoff of this potential is set at 12.0 Å.

The Lennard-Jones potential is given by:

$$U_{\mathrm{LJ}} = 4\epsilon[(\frac{\sigma}{r})^{12} - (\frac{\sigma}{r})^{6}]. \tag{S7}$$

The ion specific values of $\epsilon$ and $\sigma$ are given in Table S1, and the distance cutoff of this potential is set at 12.0 Å.

The interactions between neutral particles and ions are described by the WCA potential

$$U_{\mathrm{excl}} = \begin{cases} 4\epsilon_{\mathrm{excl}}[(\frac{\sigma}{r})^{12} - (\frac{\sigma}{r})^{6}] + \epsilon_{\mathrm{excl}} & r < r_{\mathrm{cut}} \\ 0 & r > r_{\mathrm{cut}}. \end{cases} \tag{S8}$$

$r_{\mathrm{cut}} = 2^{\frac{1}{6}}\sigma$ is located at the minimum of the corresponding Lennard-Jones potential. Values for $\sigma$ and $\epsilon_{\mathrm{excl}}$ follow the parameters given by Freeman et al.[S5] and are presented in Table S2.

### Molecular dynamics simulation details

All simulations were carried out using the software Lammps[S9] with the force fields defined in the previous two sections. Umbrella sampling simulations[S10] were performed using the Plumed software package.[S11] We used the Weighted Histogram Analysis Method (WHAM)[S12] implemented by the SMOG software package[S7] to process the simulation data and compute the free energy profiles.

#### Binding free energy of protein-DNA complexes

We carried out a series of umbrella-sampling simulations to compute the binding free energies of a set of nine protein-DNA complexes with experimentally documented binding dissociation constants.[S13–S17] Initial configurations of these simulations were prepared using the crystal structures with the corresponding PDB IDs listed in Fig. S1.

The simulations were performed under the same experimental conditions of 100 mM monovalent ions. We used a spring constant of 0.01 kcal/mol/$\text{Å}^2$ to restrain the distance between the geometric centers of protein and DNA. The centers of the umbrella windows were placed on a uniform grid of [0.0:140.0:10.0] Å, and each umbrella trajectory lasts for 7.15 million steps, with a time step of 2.0 fs. We excluded the first 3 million steps when constructing the free energy profile.

#### Single nucleosome simulations for DNA unwinding energetics

To study DNA unwinding from a 601-sequence nucleosome, we built the system by combining histone proteins with explicit coordinates for the disordered tails from PDB ID: 1KX5[S18] with the DNA structure from PDB ID: 3LZ1.[S19]

Umbrella simulations with the DNA end-to-end distance as the collective variable was performed to determine the free energy profile. The end-to-end distance was defined as the geometric center distance between the first and last five base pairs. We used a harmonic umbrella potential with a spring constant of 0.001 kcal/(mol $\cdot$ $\text{Å}^2$), and the umbrella centers were placed on a uniform grid of [30.0:510.0:30.0] Å. To increase computational efficiency, the histone core proteins and the two nucleotides located on the dyad axis of the nucleosome were rigidified during the simulations. Each umbrella trajectory lasts for 13.65 million steps, with a time step of 10 fs, and we excluded the first 3 million steps when constructing the free energy profile.

The simulation used the same ionic concentration as the experiment, which includes 0.10M NaCl and 0.5mM $MgCl_2$.[S20] The cubic simulation box size was set to 600 Å. Extra ions were added to neutralize the system. In total, the system includes a total of 13,017 $Na^+$, 65 $Mg^{2+}$, and 13003 $Cl^-$ ions.

#### Ionic dependence of the conformation for a 12-mer nucleosomal array

We constructed a nucleosomal array of 12 nucleosomes with 20-bp linker DNA to study the impact of different ions on the higher-order chromatin organization. Using the protocol outlined in a previous study,[S4] we started with a nucleosome unit extracted from the tetranucleosome crystal structure (PDB ID: 1ZBB).[S21] To connect multiple nucleosome units, we left an extra 20-bp linker DNA at the exit site of the nucleosome. We connected 12 nucleosome units to build the 12-mer nucleosomal array, with an additional 20-bp linker DNA at the end. This 20-bp extra linker of the last nucleosome unit was removed to complete the system setup. To ensure complete histone proteins with disordered tails, We replaced the histone proteins of the nucleosome units with those from the crystal structure with PDB ID: 1KX5.[S18]

To enhance conformational sampling, we performed umbrella simulations with the collective variable $Q$ defined as

$$Q = \frac{1}{N}\sum_{i}^{N}\exp(-\frac{(r_i - d_0^i)^2}{2r_0^2}). \tag{S9}$$

$i$ enumerates all the nucleosome pairs in the system and $r_i$ is the distance between the $i$-th pair. N = 66 is the number of nucleosome pairs, and $r_0$ = 20.0 Å. $d_0^i$ corresponds to the distance between the $i$-th pair of nucleosomes determined from the reference two-start structure. $Q$ measures the similarity of a given 12-mer configuration to the reference two-start structure, with larger values representing higher similarity. The reference two-start fibril structure was built in our previous study[S4] by aligning the structure with a template generated by the software fiberModel.[S22] We placed the umbrella centers at [0.40:0.90,0.1] and used a spring constant of 50.0 kcal/mol in the harmonic potentials. Each umbrella trajectory lasted 7.8 million steps, with a time step of 10 fs. We used the last 4.8 million steps to construct free energy profiles and compute ensemble averages.

We simulated the nucleosome array under four ionic conditions for comparison with experimental measurements.[S23] The 12-mer was placed in the center of a cubic box, and counter ions were added to reach the desired concentration. Excess ions were also introduced to ensure the net neutrality of the system. Specific values of the simulation box size and counter ion numbers are provided in Table S3 for reference.

#### Binding free energy between nucleosomes

##### Simulations at high salt concentrations

To determine the inter-nucleosome interactions and compare them with the DNA-origami-based experiment,[S24] we simulated a pair of 601-sequence nucleosomes. The nucleosomes were built in the same way as the single-nucleosome DNA unwrapping simulation. The simulation box size was 500 Å, and extra ions were added to neutralize the system. The system comprised a total of 2,922 $Na^+$, 828 $Mg^{2+}$, and 4,290 $Cl^-$ ions.

We defined the internal coordinate system for each nucleosome to control their relative orientations as follows. The center of each nucleosome was determined using the geometric center of the list of residues: 63-120, 165-217, 263-324, 398-462, 550-607, 652-704, 750-811, and 885-949. The IDs continuously index residues from chain A to chain H of the crystal structure with PDB ID: 1KX5. To define the nucleosome plane, we chose another point based on the geometric center of the nucleosome dyad site (residue ID: 81-131, 568-618). The unit vector pointing from the center to the center is denoted as $\vec{u}$. We further introduced another unit vector $\vec{v}$ in the nucleosome plane that points from the nucleosome center to a point defined as the geometric center of residues 63-120, 165-217, 750-811, and 885-949. Finally, the unit vector perpendicular to the nucleosome plane, $\vec{w}$, is determined as the cross product $\vec{u} \times \vec{v}$. We use $\vec{w}_1$ and $\vec{w}_2$ to differentiate the two nucleosomes.

We utilized two collective variables for the system without constraints to perform the umbrella simulations. The first collective variable measures the distance $r$ between the two nucleosome centers. The second collective variable corresponds to the angle $\theta$ between the two unit vectors, $\vec{w}_1$ and $\vec{w}_2$. The umbrella centers were placed on a uniform grid of [60.0, 130.0, 10.0] Å × [0.0, 180.0, 30.0] degrees. The spring constants of the harmonic potentials are 0.01 kcal/(mol · Å$^2$) and 0.001 kcal/(mol · ◦$^2$).

For the system that mimics the DNA-origami experiments, we imposed several spatial restraints such that the nucleosomes unbind along a specific pathway (Fig. S3). As detailed below, the restraints ensure that the first nucleosome is fixed on the X-Y plane while the second nucleosome moves along an arc 15 nm away from the origin.

1. We introduced three virtual sites, denoted as $O, A$, and $B$, with Cartesian coordinates as [0, 0, 0], [150, 0, 0], and [0, 150, 0] Å, respectively. The vectors $\overrightarrow{OA}$ and $\overrightarrow{OB}$ define the X-Y plane. We further denote the centers of the two nucleosomes as $C_1$ and $C_2$.

2. The first nucleosome was restrained at Site B using a harmonic potential with a spring constant of 100 kcal/(mol · Å$^2$). In addition, to mimic its attachment to the bottom arm of the DNA origami, we forced this nucleosome to be parallel to the X-Y plane. Specifically, we restrained the angles between $\vec{w}_1$ and $\overrightarrow{OA}$ or $\overrightarrow{OB}$ to be 90°. The spring constant of these harmonic restraints was set to 100 kcal/(mol · ◦$^2$).

3. To mimic the attachment of the second nucleosome to the upper arm of the origami, we restrained the distance between $C_2$ and Site O as 150 Å with a harmonic potential. The spring constant of this potential was set to 100.0 kcal/(mol · Å$^2$). In addition, we ensured that the second nucleosome is parallel to the plane formed by the vector $\overrightarrow{OA}$ and the vector connecting Site O to $C_2$. Two harmonic potentials were applied on the angles between $\vec{w}_2$ and $\overrightarrow{OA}$ or $\overrightarrow{OC_2}$ to restrict them to 90°. The spring constant of these restraints was set to 100 kcal/(mol · ◦$^2$). We further restricted the second nucleosome to only move in the Y-Z plane by biasing the angle between $\overrightarrow{OC_2}$ and $\overrightarrow{OA}$ to 90° with a spring constant of 100 kcal/(mol · ◦$^2$).

4. Finally, we ensured that the dyad axes of the two nucleosomes in our system are at an angle of 78°, as done experimentally,[S24] by applying a harmonic potential on the angle

between the $\vec{v}$ of the two nucleosomes. The spring constant of this potential was set to 100 kcal/(mol $\cdot \circ^2$).

We used the angle $\theta$ between the two vectors $\vec{w}_1$ and $\vec{w}_2$ as the collective variable for umbrella simulations. The umbrella centers were placed on a uniform grid of [0.0, 110.0, 5.0] degrees. The spring constant of the harmonic potential was set as 0.01 kcal/(mol $\cdot \circ^2$). Each umbrella simulation lasted 13 million steps and a time step of 10 fs. The first 3 million steps of the simulation were discarded as equilibration.

**Simulations at the physiological salt concentration**

We performed a series of simulations under the physiological salt concentration, i.e., 150 mM NaCl and 2 mM $MgCl_2$ for nucleosomes with different repeat lengths and DNA sequences. The 601-sequence nucleosomes were built in the same way as the single-nucleosome simulations. To explore the effect of DNA sequences on internucleosomal interactions, we replaced the original nucleosomal DNA with poly-dA:dT and poly-dG:dC sequences.

To investigate the effect of linker DNA, we simulated nucleosomes with a repeat length of 167 bp. We added ten base pairs of poly-dA:dT sequences on each side of the existing 147 bp 601 nucleosomal DNA using the software X3DNA.[S25] Specifically, we generated an 11-base-pair linker DNA of poly-dA:dT sequence. The additional DNA base pair was created to align the linker DNA with the existing nucleosomal DNA. This alignment was performed such that this additional base pair overlapped with the nucleosomal DNA's first or last base pair, fixing the linker DNA's orientation. Finally, we deleted the additional base pair of DNA after the alignment.

Additionally, we built nucleosomes with linker histones using a recently resolved chromatosome structure through cryoelectron microscopy (cryoEM).[S26] The experimentally determined structure (PDB ID: 7K5X) includes a 197-bp 601-sequence nucleosome with the globular domain of H1.0. As the disordered regions of the linker histone were not resolved in the structure, we modeled them based on the protein sequence using the software Modeller[S27] and connected the modeled structure to the globular domain. We then replaced the histone proteins with those from the PDB ID: 1KX5 to provide explicit coordinates for the histone tails. Only the central 167 bp of DNA was kept to build a system with 10-bp linker DNA. The globular domain of H1.0 was bound to the nucleosome dyad and simulated with the histone core protein as a rigid body for computational efficiency.

The numbers of ions and box sizes in each simulation are provided in Table S3. We employed the same two collective variables as the unrestrained simulation at high salt concentrations to conduct umbrella simulations. The umbrella centers were placed on a uniform grid of [60.0, 130.0, 10.0] Å × [0.0, 180.0, 30.0] degrees. The spring constants of the harmonic potentials are 0.01 kcal/(mol·Å$^2$) and 0.001 kcal/(mol·$\circ^2$). Each simulation lasted 13 million steps with a time step of 10 fs. We excluded the first 3 million steps when constructing the free energy profiles.

### Details of simulation analysis

#### Number of ions bound to DNA and histone proteins

To calculate the number of ions bound to the nucleosomal DNA and histone proteins, we used the COORDINATIONNUMBER command available in the Plumed[S11] software package.

For example, for every $Na^+$, we computed the coordinate number as $\mathrm{CN} = \sum_i s(r_i)$, where $i$ loops over all coarse-grained DNA sites and $r_i$ is the distance between the ion and the $i$-th DNA bead. $s(r)$ is a switching function defined as

$$s(r) = \frac{1 - (\frac{r-d_0)}{r_0})^n}{1 - (\frac{r-d_0)}{r_0})^m}, \tag{S10}$$

where $d_0 = 0.0$, $r_0 = 10.0$, $n = 15$, and $m = 30$. An ion with a coordination number greater than one was considered bound to DNA. We followed the same procedure to calculate the number of ions bound to histone proteins.

The calculations were performed using the open-source, community-developed PLUMED library,[S11] version 2.4.[S11,S28]

#### Number of unwrapped DNA base pairs

We computed the number of unwrapped DNA base pairs using a similar procedure to the one used for calculating the number of bound ions.

First, we computed a coordination number for each DNA base pair to determine whether it was bound to the histone core. The coordinate number was defined as $\mathrm{CN} = \sum_i \sum_j s(r_{i,j})$, where $i$ loops over all coarse-grained sites of the corresponding DNA base pair and $j$ loops over all coarse-grained sites of the histone core. $s(r)$ is defined in Eq. S10 with $d_0 = 0.0$, $r_0 = 8.0$, $n = 15$, and $m = 30$. A DNA base pair with CN greater than one was considered bound to histone proteins. As the histone core is not a perfect cylinder, there were several continuous regions of bound DNA interspersed by regions of unbound DNA. To avoid ambiguity, we defined the wrapped base pairs, $N_{\mathrm{wrapped}}$, as those between the first and last bound base pairs. Correspondingly, the number of unwrapped base pairs was $N_{\mathrm{unwrapped}} = 147 - N_{\mathrm{wrapped}}$.

We set $r_0 = 8.0$Å when computing the switching function. At larger values for $r_0$, we found that the calculated numbers overestimate the unwrapped base pairs, as seen from visual inspection of the structures (Fig. S10).

#### Sedimentation coefficients of nucleosome arrays

We calculated the sedimentation coefficients for the 12-mer nucleosome array using the Hull-Rad method[S29] with the following equation

$$s = 10^8 \left(\frac{M - M\bar{v}\rho_{20,w}}{N_A 6\pi\eta_0 R_T}\right). \tag{S11}$$

$M$ is the molar mass of the molecule, $N_A$ is Avogadro's number, and $\bar{v}$ is the partial specific volume. $\rho_{20,w}$ is the density of water at $20°C$, and $\eta_0$ is the water viscosity at $20°C$. $R_T$

is the translational hydrodynamic radius calculated based on the convex hull of the target biomolecule.

#### Estimation of error bars

We estimated the error bars of the 12-mer simulations based on the standard deviation calculated from the probability distribution of the variables (Fig. S2), i.e.,

$$\sigma(X) = \sqrt{E[X^2] - (E[X])^2} \tag{S12}$$

where $E[X]$ is the expected value of $X$.

We divided the trajectories into three equal-length partitions for all the other simulations and computed the free energy profiles independently. The error bars were estimated as the standard deviation of the three independent estimates.

Table S1: Summary of parameters used to describe interactions between ions and charged particles. See text *Section: Coarse-grained explicit ion model* for definitions of various parameters.

| Coarse-grained pair | $\epsilon$ (kcal/mol) | $\sigma$ (Å) | $r_{m\epsilon}$ (Å) | $\sigma_\epsilon$ (Å) | $H_1$ (kcal/mol) | $r_{mh1}$ (Å) | $\sigma_{h1}$ (Å) | $H_2$ (kcal/mol) | $r_{mh2}$ (Å) | $\sigma_{h2}$ (Å) |
|---|---|---|---|---|---|---|---|---|---|---|
| P-P | 0.18379 | 6.86 | 6.86 | 0.5 | - | - | - | - | - | - |
| $Na^+$-P | 0.02510 | 4.14 | 3.44 | 1.25 | 3.15488 | 4.1 | 0.57 | 0.47801 | 6.5 | 0.4 |
| $Na^+$-$AA^{+a}$ | 0.239 | 4.065 | 3.44 | 1.25 | 3.15488 | 4.1 | 0.57 | - | - | - |
| $Na^+$-$AA^{-b}$ | 0.239 | 4.065 | 3.44 | 1.25 | 3.15488 | 4.1 | 0.57 | 0.47801 | 6.5 | 0.4 |
| $Mg^{2+}$-P | 0.1195 | 4.87 | 3.75 | 1.0 | 1.29063 | 6.1 | 0.5 | 0.97992 | 8.3 | 1.2 |
| $Mg^{2+}$-$AA^+$ | 0.239 | 3.556 | 3.75 | 1.0 | 1.29063 | 6.1 | 0.5 | - | - | - |
| $Mg^{2+}$-$AA^-$ | 0.239 | 3.556 | 3.75 | 1.0 | 1.29063 | 6.1 | 0.5 | 0.97992 | 8.3 | 1.2 |
| $Cl^-$-P | 0.08121 | 5.5425 | 4.2 | 0.5 | 0.83652 | 6.7 | 1.5 | - | - | - |
| $Cl^-$-$AA^+$ | 0.239 | 4.8725 | 4.2 | 0.5 | 0.83652 | 6.7 | 1.5 | 0.47801 | 5.6 | 0.4 |
| $Cl^-$-$AA^-$ | 0.239 | 4.8725 | 4.2 | 0.5 | 0.83652 | 6.7 | 1.5 | - | - | - |
| $Na^+$-$Na^+$ | 0.01121 | 2.43 | 2.7 | 0.57 | 0.17925 | 5.8 | 0.57 | - | - | - |
| $Na^+$-$Mg^{2+}$ | 0.04971 | 2.37 | 2.37 | 0.5 | - | - | - | - | - | - |
| $Na^+$-$Cl^-$ | 0.08387 | 3.1352 | 3.9 | 2.06 | 5.49713 | 3.3 | 0.57 | 0.47801 | 5.6 | 0.4 |
| $Mg^{2+}$-$Mg^{2+}$ | 0.89460 | 1.412 | 1.412 | 0.5 | - | - | - | - | - | - |
| $Mg^{2+}$-$Cl^-$ | 0.49737 | 4.74 | 4.48 | 0.57 | 1.09943 | 5.48 | 0.44 | 0.05975 | 8.16 | 0.35 |
| $Cl^-$-$Cl^-$ | 0.03585 | 4.045 | 4.2 | 0.56 | 0.23901 | 6.2 | 0.5 | - | - | - |

[a] positively amino-acids, [b] negatively amino-acids

Table S2: Summary of parameters used to describe the WCA interactions between ions and neutral particles. See text *Section: Coarse-grained explicit ion model* for definitions of various parameters.

| Coarse-grained pair | $\epsilon$ (kcal/mol) | $\sigma$ (Å) |
|---|---|---|
| $Na^+$-S[a] | 0.239 | 4.315 |
| $Na^+$-A[b] | 0.239 | 3.915 |
| $Na^+$-T[c] | 0.239 | 4.765 |
| $Na^+$-G[d] | 0.239 | 3.665 |
| $Na^+$-C[e] | 0.239 | 4.415 |
| $Na^+$-AA[f] | 0.239 | 4.065 |
| $Mg^{2+}$-S | 0.239 | 3.806 |
| $Mg^{2+}$-A | 0.239 | 3.406 |
| $Mg^{2+}$-T | 0.239 | 4.256 |
| $Mg^{2+}$-G | 0.239 | 3.156 |
| $Mg^{2+}$-C | 0.239 | 3.906 |
| $Mg^{2+}$-AA[f] | 0.239 | 3.556 |
| $Cl^-$-S | 0.239 | 5.1225 |
| $Cl^-$-A | 0.239 | 4.7225 |
| $Cl^-$-T | 0.239 | 5.5725 |
| $Cl^-$-G | 0.239 | 4.4725 |
| $Cl^-$-C | 0.239 | 5.2225 |
| $Cl^-$-AA[f] | 0.239 | 4.8725 |

[a] Sugar, [b] Adenine base, [c] Thymine base, [d] Guanine base, [e] Cytosine base, [f] Non-charged amino acids

Table S3: Summary of simulation setups used in this study. Additional simulation details can be found in text *Section: Molecular dynamics simulation details.*

| Studies | Box size ($nm^3$) | Number of $Na^+$ | Number of $Mg^{2+}$ | Number of $Cl^-$ |
|---|---|---|---|---|
| Single nucleosome<br>100 mM NaCl + 0.5 mM $MgCl_2$ | 216,000 | 13,017 | 65 | 13,003 |
| 12 nucleosomes<br>5 mM NaCl | 1,331,000 | 6,196 | 0 | 4,006 |
| 12 nucleosomes<br>150 mM NaCl | 216,000 | 21,695 | 0 | 19,505 |
| 12 nucleosomes<br>0.6 mM $MgCl_2$ | 3,375,000 | 0 | 2,314 | 2,438 |
| 12 nucleosomes<br>1 mM $MgCl_2$ | 3,375,000 | 0 | 3,127 | 4,064 |
| 2 147-bp 601-seq nucleosomes<br>35 mM NaCl + 11 mM $MgCl_2$ | 125,000 | 2,922 | 828 | 4,290 |
| 2 147-bp 601-seq nucleosomes<br>150 mM NaCl + 2 mM $MgCl_2$ | 216,000 | 19,505 | 260 | 19,737 |
| 2 147-bp poly-dA:dT nucleosomes<br>150 mM NaCl + 2 mM $MgCl_2$ | 216,000 | 19,505 | 260 | 19,737 |
| 2 147-bp poly-dG:dC nucleosomes<br>150 mM NaCl + 2 mM $MgCl_2$ | 216,000 | 19,505 | 260 | 19,737 |
| 2 167-bp 601-seq nucleosomes<br>150 mM NaCl + 2 mM $MgCl_2$ | 216,000 | 19,505 | 260 | 19,657 |
| 2 167-bp 601-seq nucleosomes with H1.0<br>150 mM NaCl + 2 mM $MgCl_2$ | 216,000 | 19,505 | 260 | 19,763 |

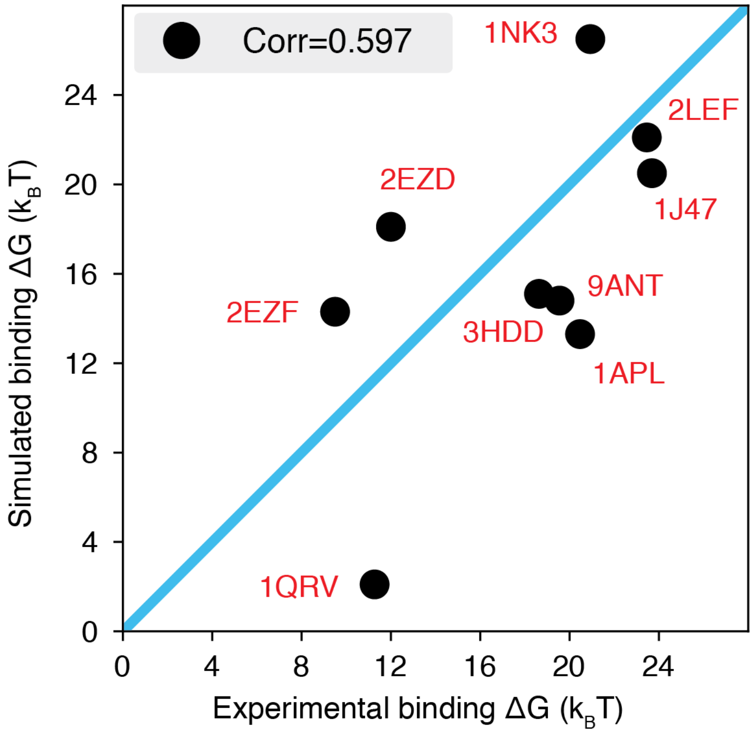


Figure S1: **The explicit-ion model predicts the binding affinities of protein-DNA complexes well, related to Fig. 1 of the main text.** Experimental and simulated binding free energies are compared for nine protein-DNA complexes,[S17] with a Pearson Correlation coefficient of 0.6. The PDB ID for each complex is indicated in red, and the diagonal line is drawn in blue. The significant correlation between simulated and experimental values supports the accuracy of the model. To further enhance the agreement between the two, it will be necessary to implement specific non-bonded interactions that can resolve differences among amino acids and nucleotides beyond simple electrostatics. Such modifications will be interesting avenues for future research. See text *Section: Binding free energy of protein-DNA complexes* for simulation details.

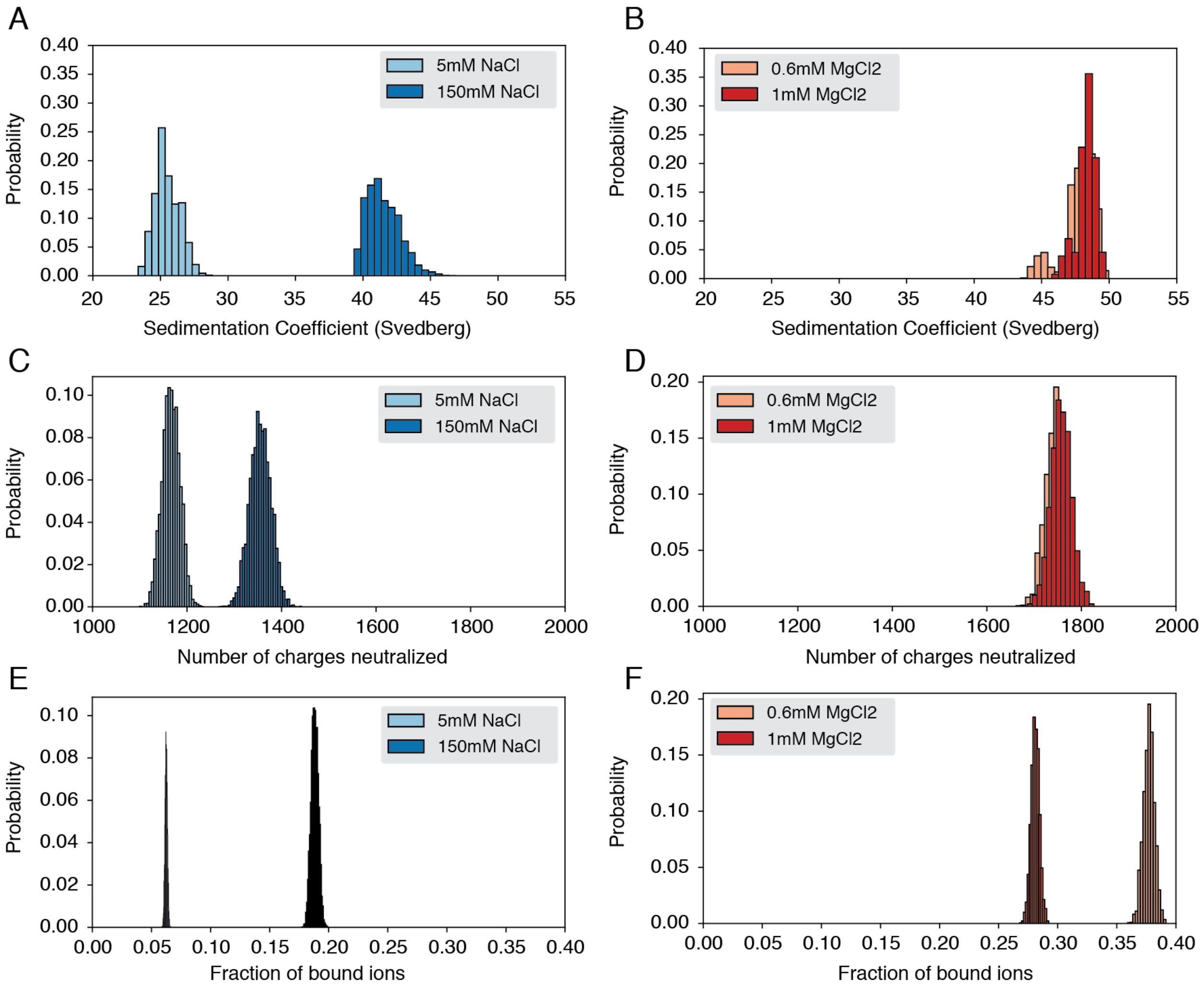


Figure S2: **Probability distributions used to compute means and standard deviations of the quantities presented in Fig. 3 of the main text.** (A) Probability distribution of sedimentation coefficients calculated from the simulation with $Na^+$ ions. (B) Probability distribution of sedimentation coefficients calculated from the simulation with $Mg^{2+}$ ions. (C) Probability distribution of neutralized charges calculated from the simulation with $Na^+$ ions. (D) Probability distribution of neutralized charges calculated from the simulation with $Mg^{2+}$ ions. (E) Probability distribution of the fraction of bound ions calculated from the simulation with $Na^+$ ions. (F) Probability distribution of the fraction of bound ions calculated from the simulation with $Mg^{2+}$ ions.

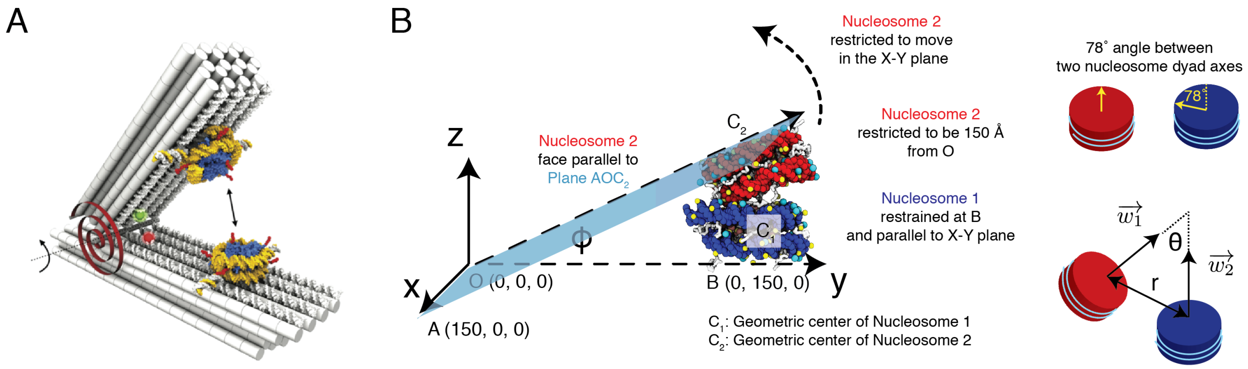


Figure S3: **Illustration of the restrained two nucleosome simulations setup, related to Fig. 4 of the main text.** (A) Schematics of the DNA-origami-based force spectrometer, reproduced from Fig. 1 of Funke et al.[S24] (B) Schematics for the spatial restraints imposed on nucleosomes in our simulations to mimic the DNA-origami setup. The vertex angle between two arms of the DNA origami system is denoted by $\Phi$. The two cartoons on the side illustrate the angle between two nucleosome dyad axes and the angle between two nucleosome planes. To define the coordinate system and other notations, please refer to *Section: Simulations at high salt concentrations* and the accompanying text.

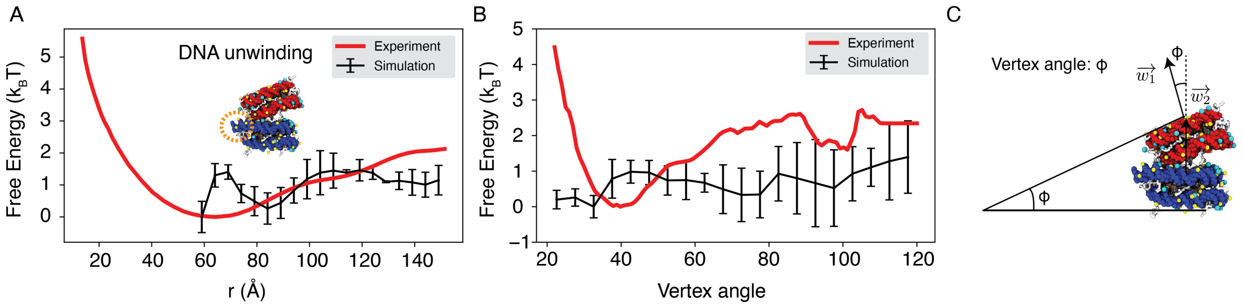


Figure S4: **Explicit ion modeling reproduces the experimental free energy profiles of nucleosome binding.** (A) Comparison between the simulated (black) and experimental (red) free energy profile as a function of the inter-nucleosome distance. Error bars were computed as the standard deviation of three independent estimates. The barrier observed between 60Å and 80Å arises from the unwinding of nucleosomal DNA when the two nucleosomes are in close proximity, as highlighted in the orange circle. (B) Comparison between the simulated (black) and experimental (red) free energy profile as a function of the vertex angle. Error bars were computed as the standard deviation of three independent estimates. (C) Illustration of the vertex angle $\Phi$ used in panel (B).

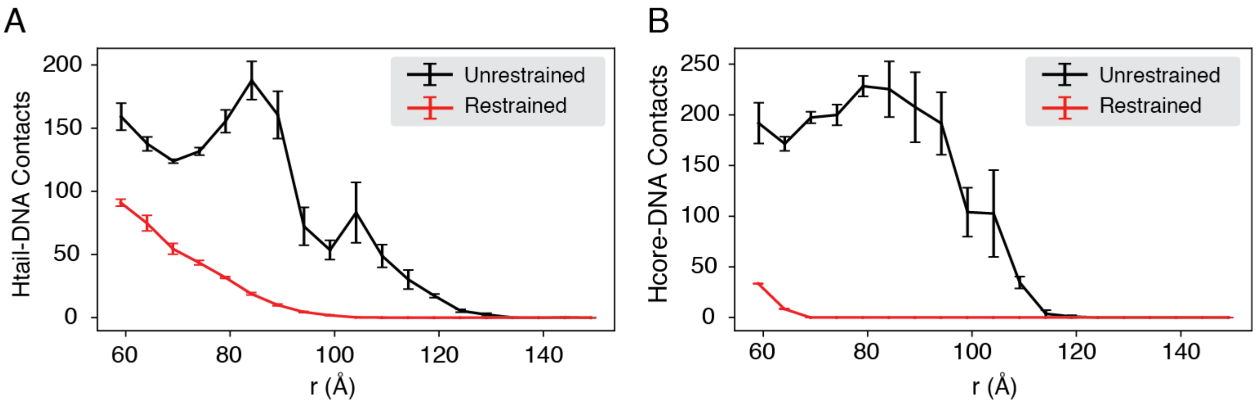


Figure S5: **Compared with DNA origami-restrained simulations, the unrestrained simulations produce more histone-DNA contacts across nucleosomes, related to Fig. 4 of the main text.** The average number of inter-nucleosome contacts between DNA and histone tails (A) or histone cores (B) are plotted as a function of the distance $r$. The error bars were estimated as the standard deviation of three equal partitions of the simulations.

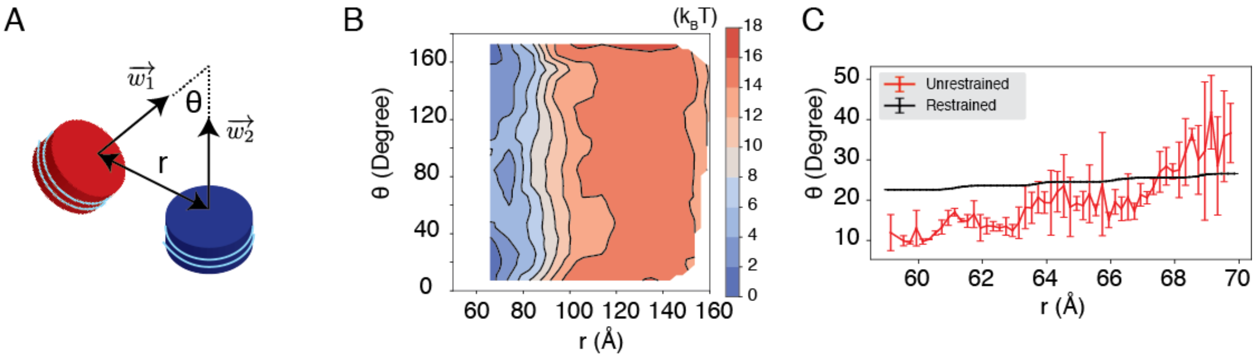


Figure S6: **The unrestricted simulations favor a smaller angle $\theta$ between two nucleosomal planes compared to the DNA origami-restrained simulations, related to Fig. 4 of the main text.** (A) Illustration of the collective variables used in the umbrella-sampling simulation. $\theta$ is the angle between two nucleosomal planes, and $r$ is the distance between the geometric centers of two nucleosomes. $\vec{w_1}$ and $\vec{w_2}$ represent the vectors perpendicular to the nucleosome planes. See text *Section: Simulations at high salt concentrations* for further definitions of the collective variables. (B) 2D Free energy landscape for nucleosome interactions under 35 mM NaCl and 11 mM $MgCl_2$ salt, plotted as a function of $r$ and $\theta$. (C) The average value of $\theta$ as a function of the distance $r$ for the unrestricted (red) and the DNA origami-restrained (black) simulations. The error bars were estimated as the standard deviation of three equal partitions of the simulations.

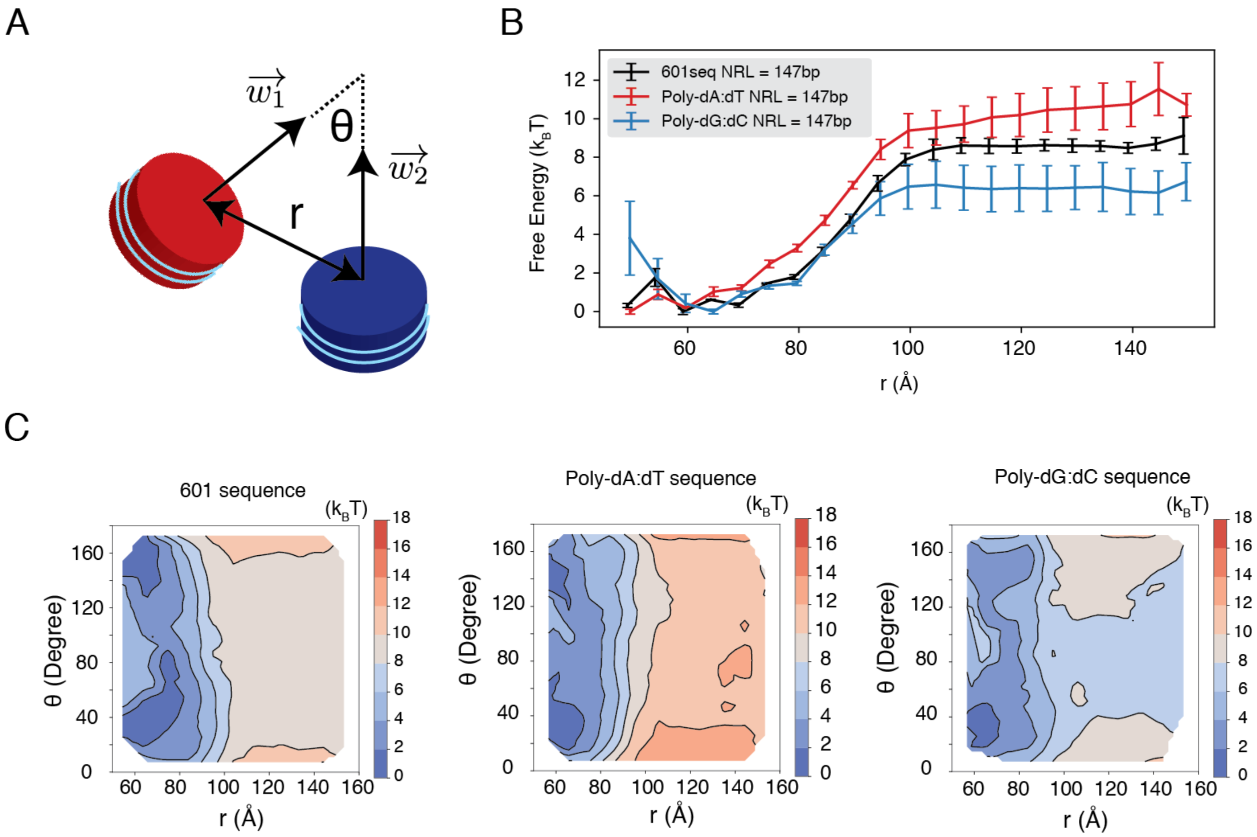


Figure S7: **Dependence of internucleosome interactions on the DNA sequence, related to Fig. 5 of the main text.** See text *Section: Simulations at the physiological salt concentration* for further discussions on simulation details. (A) Illustration of the collective variables used in umbrella-sampling simulations. $\theta$ is the angle between two nucleosomal planes, and $r$ is the distance between the geometric centers of two nucleosomes. $\vec{w_1}$ and $\vec{w_2}$ represent the vectors perpendicular to the nucleosome planes. (B) The free energy profile as a function of the distance $r$ between the geometric centers of two nucleosomes with 601, poly-dA:dT, and poly-dG:dC sequences. (C) The 2D free energy profiles as a function of $\theta$ and $r$. The simulations used nucleosomes with 601, poly-dA:dT, and poly-dG:dC sequences.

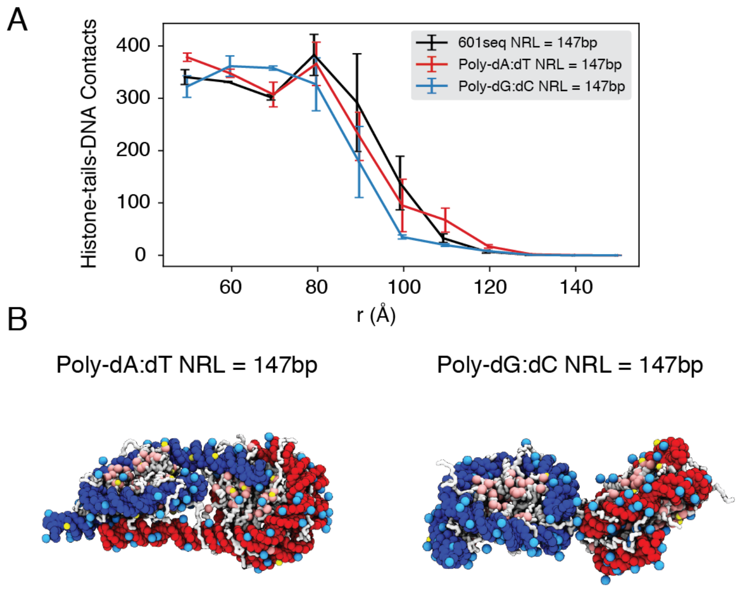


Figure S8: **The poly-dA:dT sequence produces a higher number of cross-nucleosome histone-DNA contacts compared to the poly-dG:dC sequence, related to Fig. 5 of the main text.** (A) The average number of inter-nucleosome contacts between histone proteins and nucleosomal DNA is plotted as a function of the distance $r$ between the geometric centers of two nucleosomes. The error bars are estimated as the standard deviation of three equal partitions of the simulations. (B) Representative structures from simulations with poly-dA:dT (left) and poly-dG:dC (right) nucleosomes. Noticeable DNA unwrapping can be seen in poly-dA:dT nucleosomes, contributing to the increased cross-nucleosome contacts.

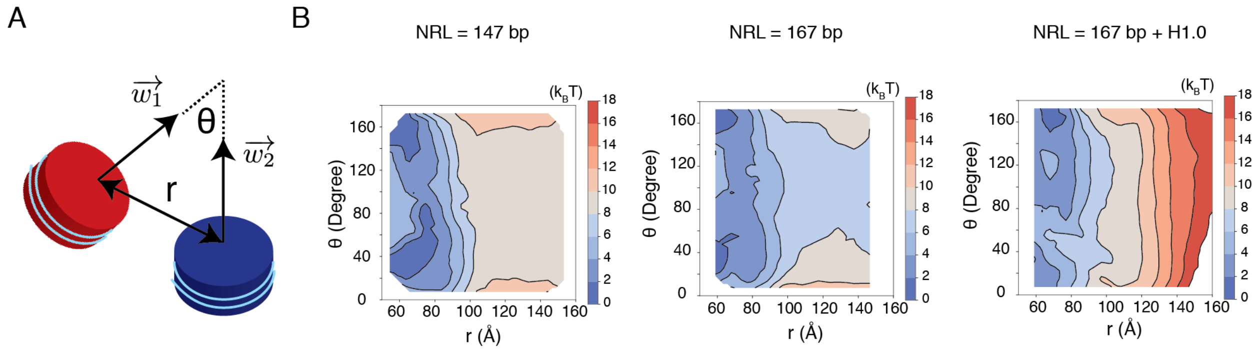


Figure S9: **Free energy profiles for the interactions between a pair of nucleosomes at different nucleosome repeat lengths (NRL) and in the presence of the linker histone H1.0, related to Fig. 5 of the main text.** See text *Section: Simulations at the physiological salt concentration* for further discussions on simulation details. (A) Illustration of the collective variables used in the umbrella-sampling simulations. $\theta$ is the angle between two nucleosome planes, and $r$ is the distance between the geometric centers of two nucleosomes. $\vec{w_1}$ and $\vec{w_2}$ represent the vectors perpendicular to the nucleosome planes. (B) 2D free energy profiles as a function of $\theta$ and $r$ for the three systems indicated in the titles.

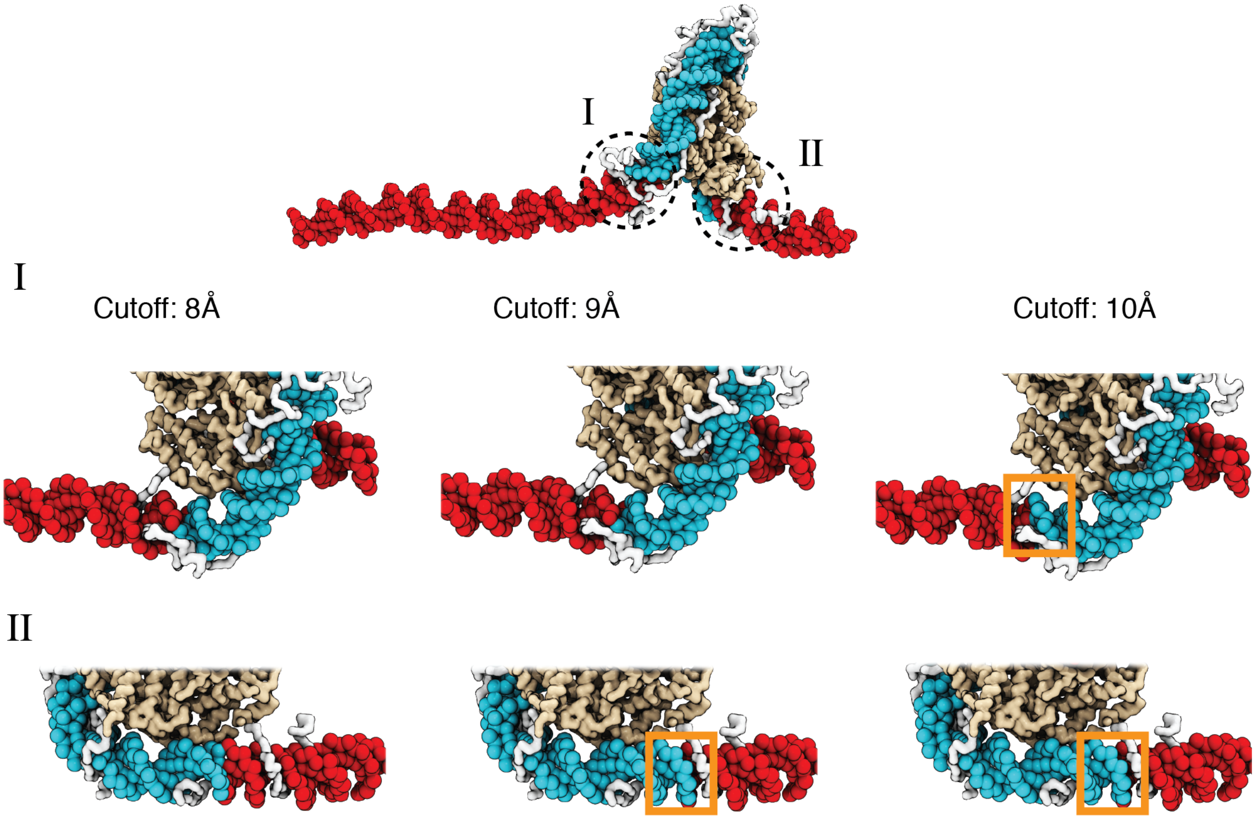


Figure S10: **A cutoff value of 8.0 Å produces more accurate values for the number of unwrapped DNA base pairs as determined from visual inspection of representative configurations, related to Fig. 2 of the main text.** See text *Section: Number of unwrapped DNA base pairs* for additional discussions. A typical nucleosome structure with most of the outer layer DNA unwrapped was used to examine the impact of different cutoff values. The histone core is colored in gold, with histone tails in white, the wrapped DNA in blue, and the unwrapped DNA in red. The discrepancy among various cutoff values is evident in the highlight regions enclosed by dotted circles. As shown in the zoom-ins in the middle panel, a cutoff of 10 Å results in 3 additional base pairs of DNA detected as wrapped in *I* (highlighted in orange square). However, these extra base pairs not detected with a cutoff of 8 Å are visibly detached from histone proteins. Similarly, 9 Å and 10 Å cutoff values result in 5 extra base pairs of DNA detected as wrapped in *II* (highlighted in orange squares in the bottom panel).

(S1) Freeman, G. S.; Hinckley, D. M.; Lequieu, J. P.; Whitmer, J. K.; de Pablo, J. J. Coarse-grained modeling of DNA curvature. *The Journal of Chemical Physics* **2014**, *141*, 165103.

(S2) Zhang, B.; Zheng, W.; Papoian, G. A.; Wolynes, P. G. Exploring the Free Energy Landscape of Nucleosomes. *J. Am. Chem. Soc.* **2016**, *138*, 8126–8133.

(S3) Ding, X.; Lin, X.; Zhang, B. Stability and folding pathways of tetra-nucleosome from six-dimensional free energy surface. *Nat Commun* **2021**, *12*, 1091.

(S4) Liu, S.; Lin, X.; Zhang, B. Chromatin fiber breaks into clutches under tension and crowding. *Nucleic Acids Research* **2022**, *50*, 9738–9747.

(S5) Freeman, G. S.; Hinckley, D. M.; de Pablo, J. J. A coarse-grain three-site-per-nucleotide model for DNA with explicit ions. *The Journal of Chemical Physics* **2011**, *135*, 165104.

(S6) Noel, J. K.; Whitford, P. C.; Onuchic, J. N. The Shadow Map: A General Contact Definition for Capturing the Dynamics of Biomolecular Folding and Function. *J. Phys. Chem. B* **2012**, *116*, 8692–8702.

(S7) Noel, J. K.; Levi, M.; Raghunathan, M.; Lammert, H.; Hayes, R. L.; Onuchic, J. N.; Whitford, P. C. SMOG 2: A Versatile Software Package for Generating Structure-Based Models. *PLoS Comput Biol* **2016**, *12*, e1004794.

(S8) Miyazawa, S.; Jernigan, R. L. Estimation of effective interresidue contact energies from protein crystal structures: quasi-chemical approximation. *Macromolecules* **1985**, *18*, 534–552.

(S9) Plimpton, S. Fast Parallel Algorithms for Short-Range Molecular Dynamics. *Journal of Computational Physics* **1995**, *117*, 1–19.

(S10) Torrie, G.; Valleau, J. Nonphysical sampling distributions in Monte Carlo free-energy estimation: Umbrella sampling. *Journal of Computational Physics* **1977**, *23*, 187–199.

(S11) The PLUMED consortium, Promoting transparency and reproducibility in enhanced molecular simulations. *Nat Methods* **2019**, *16*, 670–673.

(S12) Kumar, S.; Rosenberg, J. M.; Bouzida, D.; Swendsen, R. H.; Kollman, P. A. THE weighted histogram analysis method for free-energy calculations on biomolecules. I. The method. *J. Comput. Chem.* **1992**, *13*, 1011–1021.

(S13) Dragan, A. I.; Klass, J.; Read, C.; Churchill, M. E.; Crane-Robinson, C.; Privalov, P. L. DNA Binding of a Non-sequence-specific HMG-D Protein is Entropy Driven with a Substantial Non-electrostatic Contribution. *Journal of Molecular Biology* **2003**, *331*, 795–813.

(S14) Dragan, A. I.; Read, C. M.; Makeyeva, E. N.; Milgotina, E. I.; Churchill, M. E.; Crane-Robinson, C.; Privalov, P. L. DNA Binding and Bending by HMG Boxes: Energetic Determinants of Specificity. *Journal of Molecular Biology* **2004**, *343*, 371–393.

(S15) Dragan, A. I.; Liggins, J. R.; Crane-Robinson, C.; Privalov, P. L. The Energetics of Specific Binding of AT-hooks from HMGA1 to Target DNA. *Journal of Molecular Biology* **2003**, *327*, 393–411.

(S16) Dragan, A. I.; Li, Z.; Makeyeva, E. N.; Milgotina, E. I.; Liu, Y.; Crane-Robinson, C.; Privalov, P. L. Forces Driving the Binding of Homeodomains to DNA. *Biochemistry* **2006**, *45*, 141–151.

(S17) Privalov, P. L.; Dragan, A. I.; Crane-Robinson, C. Interpreting protein/DNA interactions: distinguishing specific from non-specific and electrostatic from non-electrostatic components. *Nucleic Acids Research* **2011**, *39*, 2483–2491.

(S18) Davey, C. A.; Sargent, D. F.; Luger, K.; Maeder, A. W.; Richmond, T. J. Solvent Mediated Interactions in the Structure of the Nucleosome Core Particle at 1.9Å Resolution. *Journal of Molecular Biology* **2002**, *319*, 1097–1113.

(S19) Vasudevan, D.; Chua, E. Y. D.; Davey, C. A. Crystal structures of nucleosome core particles containing the '601' strong positioning sequence. *J Mol Biol* **2010**, *403*, 1–10.

(S20) Hall, M. A.; Shundrovsky, A.; Bai, L.; Fulbright, R. M.; Lis, J. T.; Wang, M. D. High-resolution dynamic mapping of histone-DNA interactions in a nucleosome. *Nat Struct Mol Biol* **2009**, *16*, 124–129.

(S21) Schalch, T.; Duda, S.; Sargent, D. F.; Richmond, T. J. X-ray structure of a tetranucleosome and its implications for the chromatin fibre. *Nature* **2005**, *436*, 138–141.

(S22) Koslover, E. F.; Fuller, C. J.; Straight, A. F.; Spakowitz, A. J. Local Geometry and Elasticity in Compact Chromatin Structure. *Biophysical Journal* **2010**, *99*, 3941–3950.

(S23) Correll, S. J.; Schubert, M. H.; Grigoryev, S. A. Short nucleosome repeats impose rotational modulations on chromatin fibre folding. *EMBO J* **2012**, *31*, 2416–2426.

(S24) Funke, J. J.; Ketterer, P.; Lieleg, C.; Schunter, S.; Korber, P.; Dietz, H. Uncovering the forces between nucleosomes using DNA origami. *Sci. Adv.* **2016**, *2*, e1600974.

(S25) Lu, X.-J.; Olson, W. K. 3DNA: a software package for the analysis, rebuilding and visualization of three-dimensional nucleic acid structures. *Nucleic Acids Res* **2003**, *31*, 5108–5121.

(S26) Zhou, B.-R.; Feng, H.; Kale, S.; Fox, T.; Khant, H.; De Val, N.; Ghirlando, R.; Panchenko, A. R.; Bai, Y. Distinct Structures and Dynamics of Chromatosomes with Different Human Linker Histone Isoforms. *Molecular Cell* **2021**, *81*, 166–182.e6.

(S27) Eswar, N.; Webb, B.; Marti-Renom, M. A.; Madhusudhan, M.; Eramian, D.; Shen, M.; Pieper, U.; Sali, A. Comparative Protein Structure Modeling Using Modeller. *CP in Bioinformatics* **2006**, *15*.

(S28) Tribello, G. A.; Bonomi, M.; Branduardi, D.; Camilloni, C.; Bussi, G. PLUMED 2: New feathers for an old bird. *Computer Physics Communications* **2014**, *185*, 604–613.

(S29) Fleming, P. J.; Fleming, K. G. HullRad: Fast Calculations of Folded and Disordered Protein and Nucleic Acid Hydrodynamic Properties. *Biophysical Journal* **2018**, *114*, 856–869.
